# Potassium homeostasis and signaling as a determinant of *Echinacea* species tolerance to salinity stress

**DOI:** 10.1101/2022.10.10.511607

**Authors:** Fatemeh Ahmadi, Abbas Samadi, Ebrahim Sepehr, Amir Rahimi, Sergey Shabala

## Abstract

Salt tolerant is strongly related to potassium (K^+^) retention in plant tissues under salt stress conditions. However, it is unclear for different *Echinacea* species. So, mechanistic basis of four *Echinacea* species (i.e. *Echinacea purpurea, Echinacea angustifolia, Echinacea pallida*, and *Echinacea sanguinea*) to salinity stress tolerance, and K^+^ retention were assessed in the present study. Non-invasive microelectrode ion flux measuring, DHAR and MDHAR activities, and pharmacological measurements were performed based on the standard methods. Ion flux measurements revealed higher K^+^ efflux in *E. pallida* and *E. sanguinea* species compared to the *E. purpurea* and *E. angustifolia* species in the elongation zone. Higher salinity-induced H^+^ efflux was found in the elongation zone than mature zone. However, *E. angustifolia* and *E. purpurea* had more Ca^2+^ influx compared to *E. pallida* and *E. sanguinea* species. Net K^+^ efflux decreased (> 90%) in the presence of TEA and GdCl_3_. Increasing of Ca^2+^ uptake and K^+^ loss in four *Echinacea* species roots were found in the presence of 0.3 mM Cu/Ascorbate (Cu/Asc). The significant role of H^+^-ATPase in H^+^ efflux was demonstrated by Sodium orthovanadate. Ultimately, the physiological properties of *Echinacea* species have a critical role in salinity-resistant/sensitive differences. Future scientific understanding of *Echinacea* species physiognomies may be necessary for better understanding of the plant behavior to salinity stress.

**One-sentence summary:** Higher K^+^ efflux in *E. pallida* and *E. sanguinea* species as a result of NaCl and ROS act as a metabolic switch to save energy for adaptations and repairs in salinity stress conditions.

## INTRODUCTION

The importance of potassium (K^+^) as an essential macronutrient in plants and a rate-limiting element for enzymatic activities is well demonstrated (Ma et al., 2022). El-Mageed et al., (2022) reported that the plant’s reaction to different abiotic stresses, especially salinity, is significantly affected by K^+^ content. Meanwhile, the key role of K^+^ in protein synthesis (Abdelraouf et al., 2022) and phloem sugar loading (Mondal et al., 2022) is well recognized. The critical reactions such as photosynthesis and osmotic adjustment were disturbed in K^+^-deficiency plants (Feghhenabi et al., 2022). Signaling reaction of K^+^ has been recognized (Rashidi et al., 2022).

Salinity stress, as worldwide abiotic stress, cause to increase plant’s chronic K^+^ deficiency, while the plants can utilize the salinity stress by adequate available K^+^ (Dave et al., 2022). Potassium loss in the presence of NaCl is a usual phenomenon, and the salt-sensitive species were more affected than tolerant (Khan et al., 2022). Different pathways are recognized in both sensitive and tolerant plant species to salinity stress. It is reported (Francini et al., 2022) that guard cell outward rectifying K^+^ channel (GORK) and nonselective cation channels (ROS-activated NSCC channels) are responsible for mediating K^+^ efflux. Shabala et al., (2022) reported that the effective role of plasma membrane potential on GORK channels activation and K^+^ efflux as affected by salinity stress. Tolerance plants to salinity stress are able to retention K^+^ in plants cell, which is demonstrated in wheat (Cuin et al., 2012), barley (Wu et al., 2015b), maize (Gao et al., 2016), and oilseed crops (Chakraborty et al., 2016). The ATP production is limited in salt-affected plants (Wang et al., 2022), and they spend most of ATP on defense purposes (Chattha et al., 2022). Thus, the plant’s defense ability is strongly dependent on metabolic activity.

In this condition, the metabolic switch of K^+^ efflux is suggested, causing to decrease in the energy-consuming biosynthesis and consumption of more ATP for defense purposes (Han et al., 2022). So, how do plant species cope with this dilemma? Is maintaining the K^+^ unchanged better than the K^+^ loss? Which pathway will result in higher salt affected-plants productivity?

The best way to answer these questions is to pay attention to plant strategies carefully. It is reported that Na^+^ exclusion is an important mechanism under salinity stress conditions and is controlled by genetics (Shahzad et al., 2022). The gene can encode the Na^+^/H^+^ exchanger in the plasma membrane (PM) and utilize the exceeded Na^+^ (Shabala et al., 2022). However, it requires H^+^-ATPase operation to osmotic adjustment, as an energy-consuming process (El-Mageed et al., 2022). On the other hand, numerous roots and shoot reactive oxygen species (ROS), especially hydrogen peroxide (H_2_O_2_), were produced as affected by salinity stress in plants and result in oxidative stress (Miller et al., 2010). However, previous researches demonstrated that different ions permeable channels were activated as affected by ROS (Shabala et al., 2016). So, how is this critical pathway resolved by different *Echinacea* species, such as *Echinacea purpurea, Echinacea angustifolia, Echinacea pallida*, and *Echinacea sanguinea*? What are the differences between *Echinacea* species reactions to salinity stress? Which species are most tolerant to salinity stress and which one is more sensitive?

To answer the above gaps, four *Echinacea* species, including *E. purpurea, E. angustifolia, E. pallida*, and *E. sanguinea*, were studied electro physiologically based on ions flux measurements, evaluation of DHAR and MDHAR activities, and pharmacological studies in the both elongation and mature root zones.

## RESULTS

### K^+^ flux as affected by NaCl stress

The K^+^ efflux in elongation and mature root zones of four *Echinacea* species is shown in **Figure 1(A-D)**. Based on the results, salinity stress (100 mM NaCl) could affect the net K^+^ efflux significantly in elongation and mature root zones **(Figure 1 A-B)**. As shown in **Figure 1C**, net K^+^ efflux was three times higher in the root elongation zone and caused the main difference in K^+^ loss between elongation and mature root zones. Potassium efflux in *E. pallida* and *E. sanguinea* species were higher than *in E. purpurea* and *E. angustifolia* in the elongation zone, relevant that the signaling role of K^+^ efflux in *E. pallida* and *E. sanguinea* species is possible. The salt-tolerant *E. angustifolia* had better K^+^ retention in comparison with salt-sensitive *E. purpurea* in the root mature zone **(Figure 1C)**. According to pharmacological results, TEA and GdCl_3_ blockers remarkably reduced (70% inhabitation, **Figure 1D**) the salt-induced K^+^ efflux, demonstrating the important positive role of GORK and NSCC channels to control the NaCl--induced K^+^ efflux.

**Figure 1.**
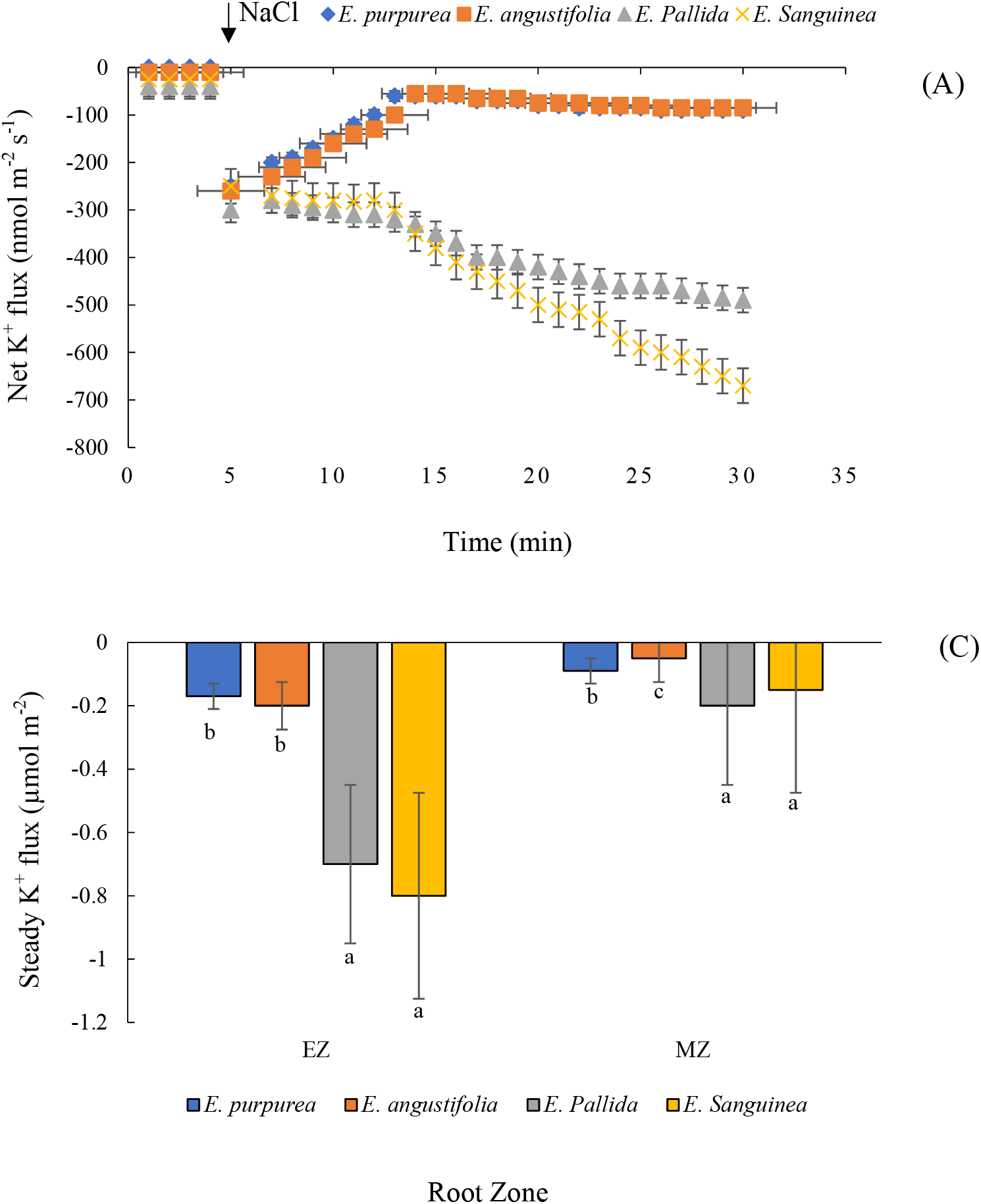
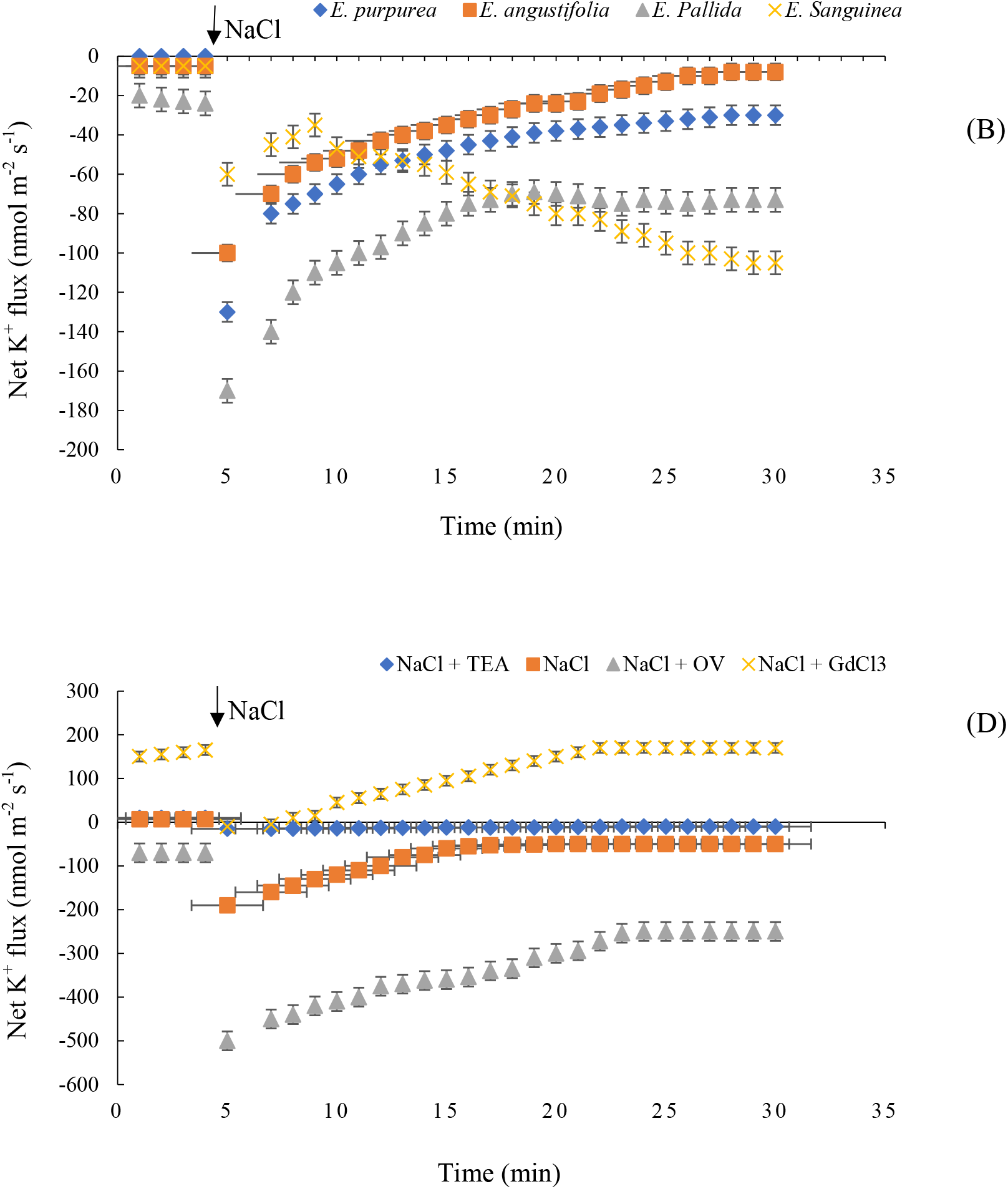
Transiently measured net K^+^ flux responses of four *Echinacea* species measured from root elongation (EZ; A) and mature (MZ; B) zones exposed to 100 mM NaCl stress. Total K^+^ fluxes (C) measured from EZ and MZ were calculated over 20 min after the addition of 100 mM NaCl. Transient net K^+^ flux response (D) to 100 mM NaCl measured from EZ in the presence of ion transport inhibitors: 1 mM sodium orthovanadate (OV), plasma membrane (PM) H^+^-ATPase blocker; 20 mM tetraethylammonium chloride (TEA), K^+^-selective PM channels blocker; 0.1 mM gadolinium chloride (GdCl_3_), known as non-selective cation channel (NSCC) blocker. Different letters indicate the significant difference at * P < 0.05 among different *Echinacea* species. The error bars indicate the standard error (SE) for all the replicated data for each treatment. Data are shown as mean ± SE (n = 5).

### NaCl-induced H^+^ flux response in roots

Obtained results demonstrated the increase of H^+^ efflux in the presence of 100 mM NaCl in all *Echinacea* species (**Figure 2 AB)**, which was higher in the root elongation zone than mature zone. The highest H^+^ efflux was found in *E. angustifolia* species, however, the least H^+^ efflux was found in *E. pallida* and *E. sanguinea* species, suggesting the different strategies of these species to resolve the salinity stress. Based on the H^+^ efflux kinetics **(Figure 2C)**, strong species specificity in different *Echinacea* species are as *E. angustifolia* > *E. purpurea* > *E. sanguinea* > *E. pallida*. It is ∼2–3-fold higher in *E. angustifolia* than *E. purpurea* in the root elongation zone **(Fig. 2C)**. According to pharmacological results **(Figure 2D)**, 1 Mm OV could decrease the salt-induced H^+^ efflux more than 80%, demonstrating the role of H^+^-ATPase in NaCl-induced H^+^ efflux. So, it can be considered that the activation of H^+^-ATPase as affected by salinity stress is not probable in the *E. angustifolia* and *E. purpurea* species.

**Figure 2.**
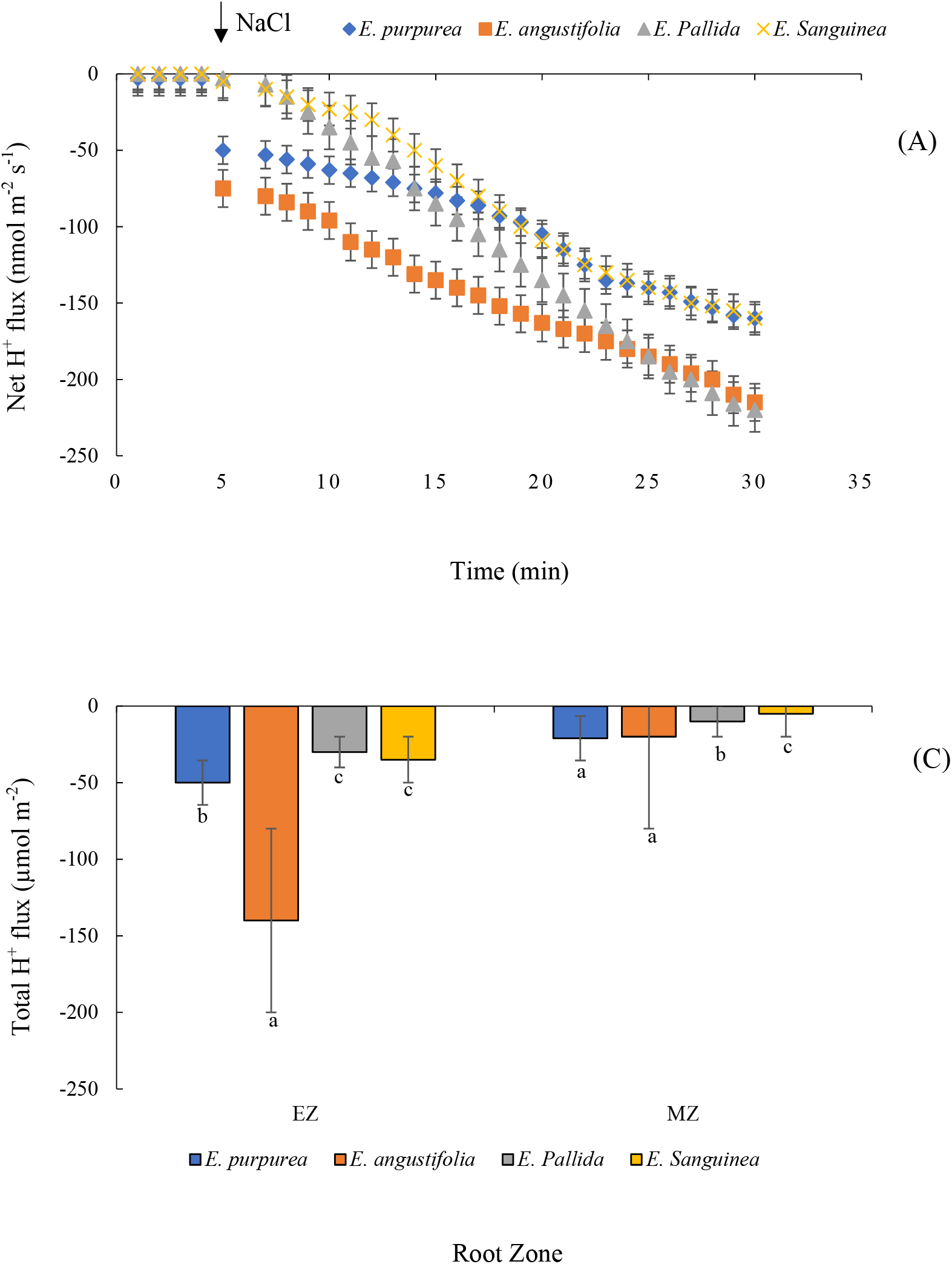
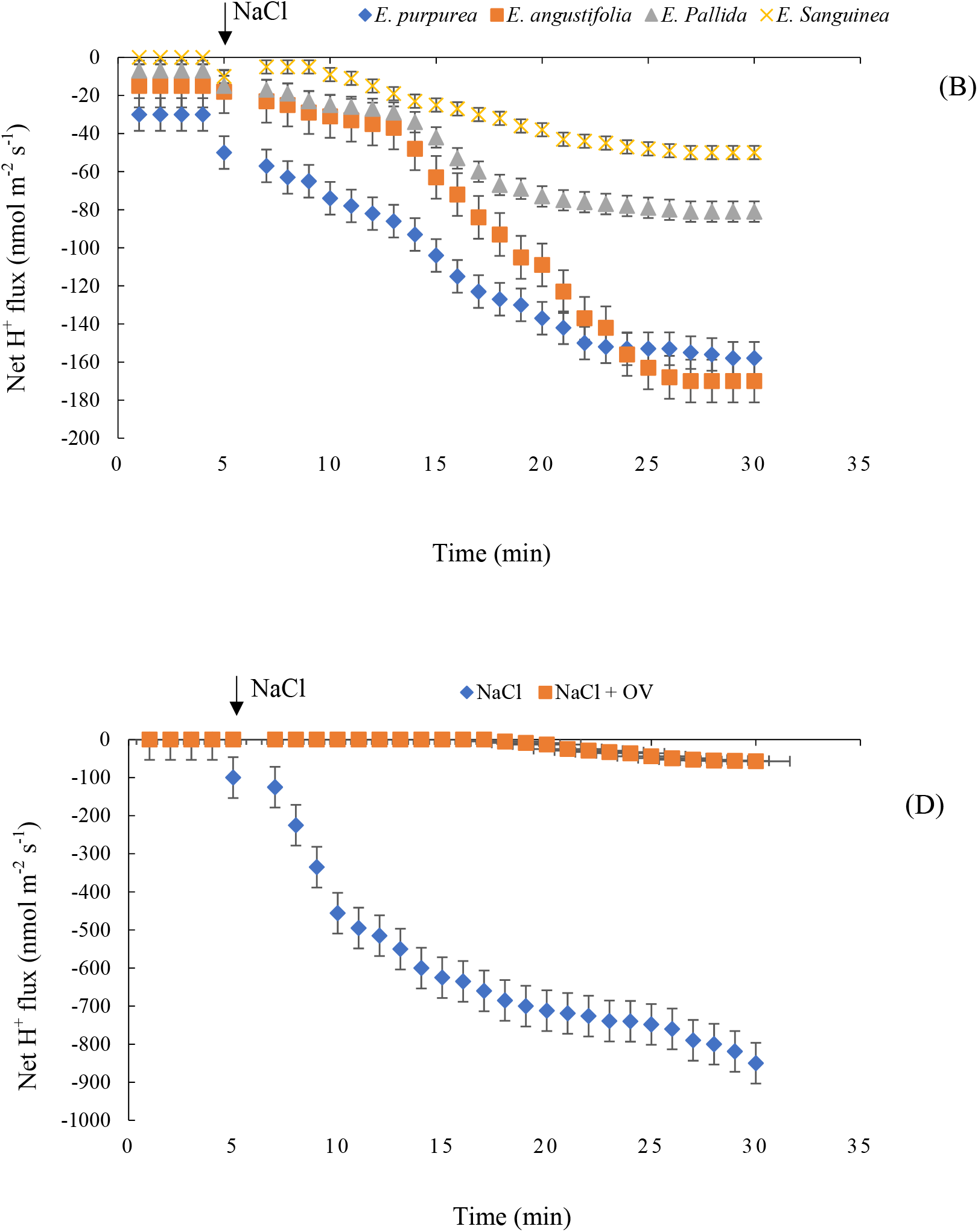
Transiently measured net H^+^ flux responses of four *Echinacea* species measured from root elongation (EZ; A) and mature (MZ; B) zones exposed to 100 mM NaCl stress. Total H^+^ flux (C) measured from EZ and MZ was calculated over 20 min after the addition of 100 mM NaCl. Transient net H^+^ flux response (D) to 100 mM NaCl was measured from EZ in the presence of a plasma membrane (PM) H^+^-ATPase blocker sodium orthovanadate (OV; 1 mM). Different letters indicate the significant difference at * P < 0.05 among different *Echinacea* species. The error bars indicate the standard error (SE) for all the replicated data for each treatment. Data are shown as mean ± SE (n = 5).

### Ions flux as affected by ROS

Salinity stress has a significant effect on ROS accumulation in the root system (Bose et al., 2014). It is reported that the sensitivity of ion channels to ROS could affect the plant’s salt tolerance (Bose et al., 2014). The H_2_O_2_-induced K^+^ fluxes in both elongation and mature root zones are shown in **Figure 3AB**. Accordingly, a massive K^+^ efflux was found in all *Echinacea* species in the presence of 10 mM H_2_O_2_. Based on the results, the highest peak K^+^ efflux was found in salt-tolerant *E. pallida* and *E. angustifolia* species **(Figure 3AB)**. No clear pattern was found for the mature zone. Total K^+^ loss in *E. pallida* and *E. angustifolia* species was ∼3-fold higher than in other species, relevant that H_2_O_2_-induced K^+^ efflux act as a signal intolerant species **(Fig. 3C)**. The net K^+^ efflux was more than 90% lower in the presence of TEA and GdCl_3_ blockers than 10 mM H_2_O_2_ only **(Figure 3D)**. No K^+^ efflux inhabitation was found with DPI, a blocker of NADPH oxidase **(Figure 3D)**.

**Figure 3.**
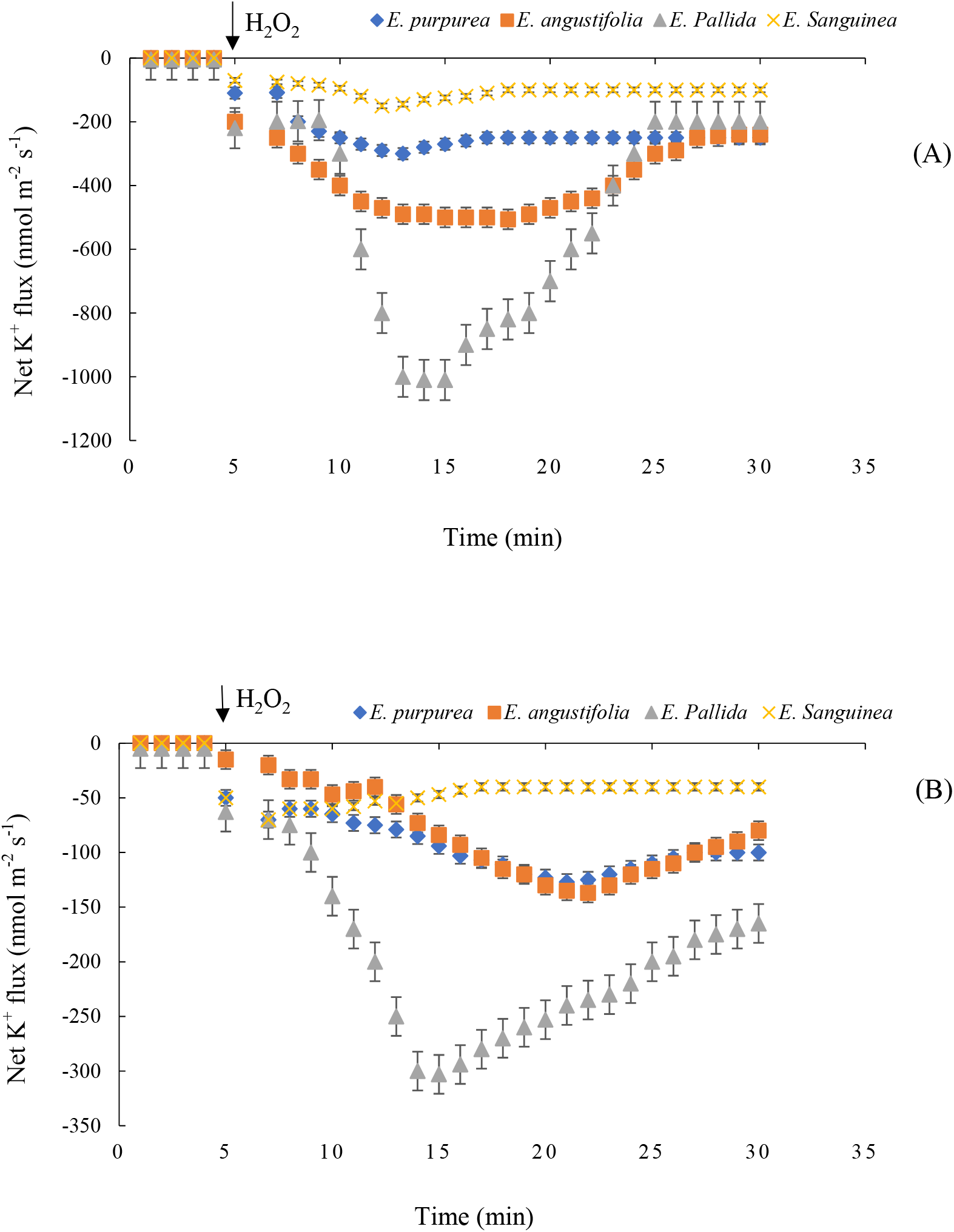
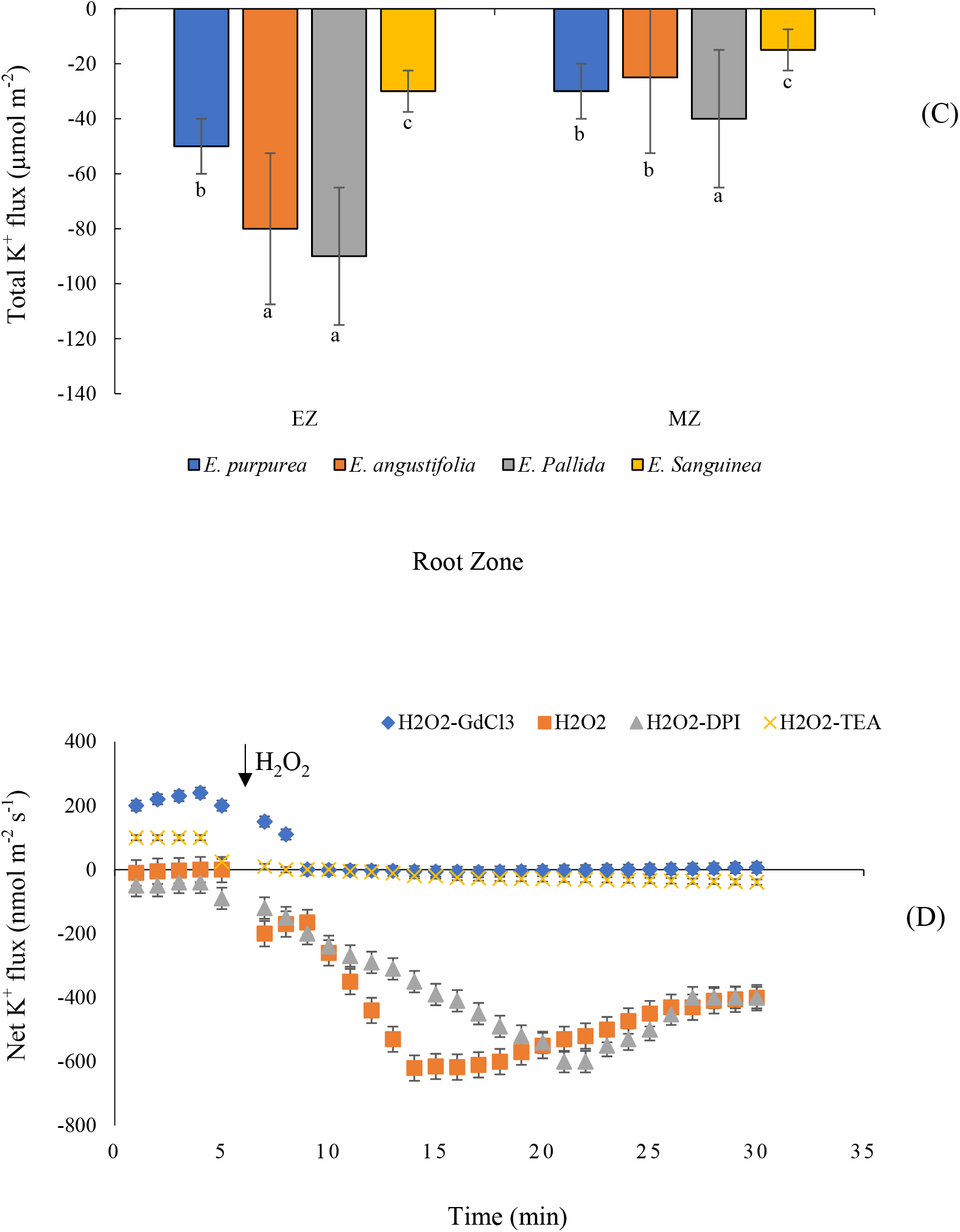
Transiently measured net K+ flux responses of four *Echinacea* species measured from root elongation (EZ; A) and mature (MZ; B) zones exposed to 10 mM H_2_O_2_ stress. Total K^+^ efflux (C) measured from EZ and MZ was calculated over 20 min after the addition of 10 mM H_2_O_2_ stress. Transient net K^+^ flux (D) response to 10 mM H_2_O_2_ stress measured from EZ for 1 h with inhibitors: 20 µM diphenyleneiodonium (DPI), NADPH oxidase blocker; 20 mM tetraethylammonium chloride (TEA), K^+^-selective PM channels blocker; 0.1 mM gadolinium chloride (GdCl_3_), known as non-selective cation channel (NSCC) blocker. Different letters indicate the significant difference at * P < 0.05 among different *Echinacea* species. The error bars indicate the standard error (SE) for all the replicated data for each treatment. Data are shown as mean ± SE (n = 5).

### Calcium flux as affected by H_2_O_2_

Potassium efflux across the plasma membrane is related to the Ca^2+^ influx in the root, and a negative correlation is present between salinity tolerance and ROS-stimulated Ca^2+^ uptake in plants (Wang et al., 2018). Accordingly, the plasma membrane Ca^2+^-permeable channel’s sensitivity to 10 mM H_2_O_2_ was assessed in four *Echinacea* species **(Figure 4AB)**. A massive Ca^2+^ influx was obtained in the presence of 10 mM H_2_O_2_ in all *Echinacea* species in both root zones which was much stronger in tolerant *Echinacea* species **(Figure 4AB)**. The higher Ca^2+^ influx peak was found in the *E. angustifolia* species in the elongation zone (**Figure 4C)**. The highest peak of Ca^2+^ influx in a mature zone was found in *E. angustifolia* and *E. purpurea* species **(Figure 4C)**. The magnitude of both peak Ca^2+^ uptake and the overall amount of Ca^2+^ taken was ranked as *E. angustifolia* > *E. purpurea* > *E. pallida* > *E. sanguinea* **(Figure 4C)**. The Ca^2+^ influx was decreased by more than 90% in the presence of 0.1 mM of GdCl_3_; however, it was inhabited by more than 50% in the presence of 20 µM DPI under 10 mM H_2_O_2_ stress **(Figure 4D)**.

**Figure 4.**
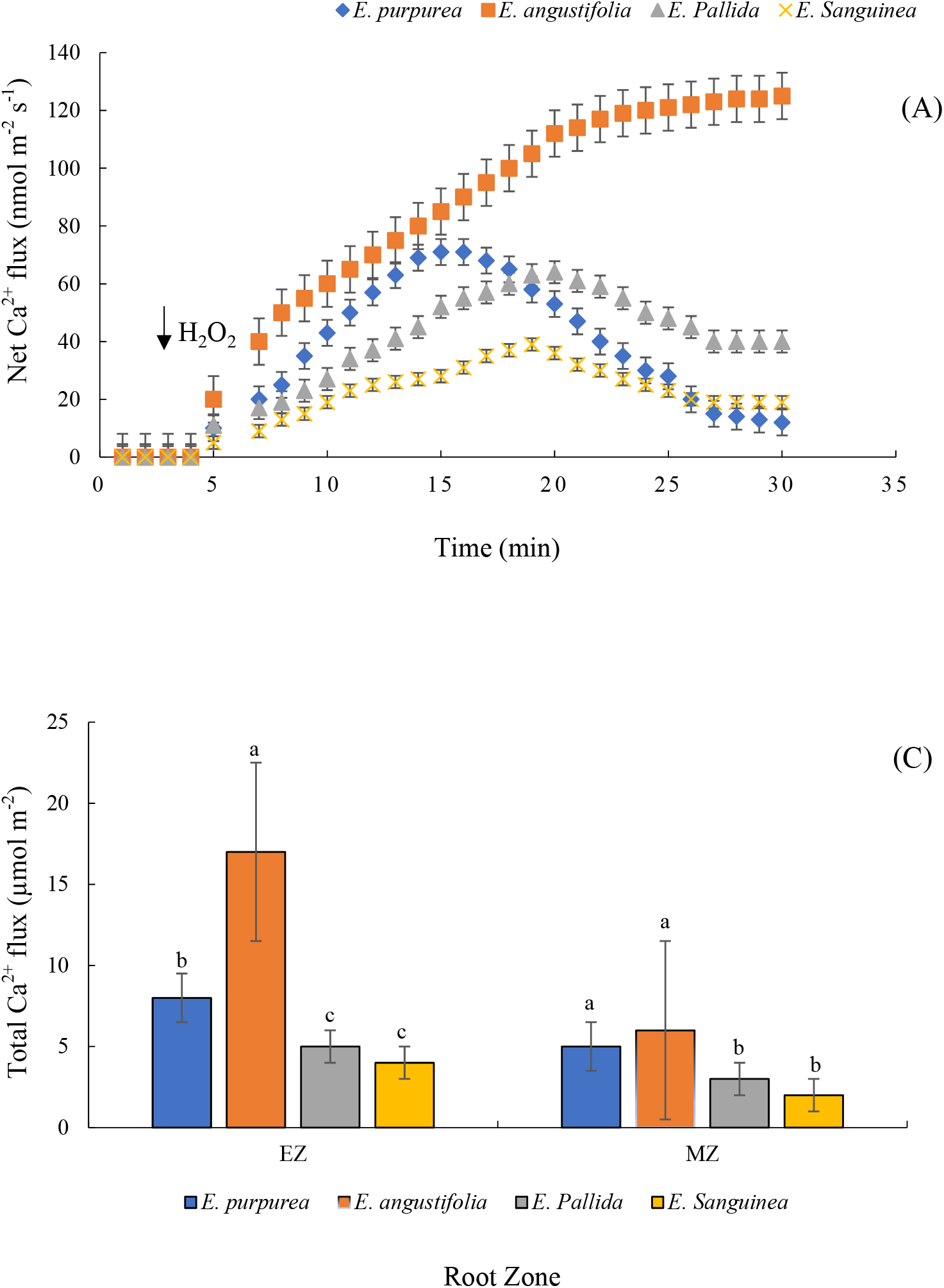
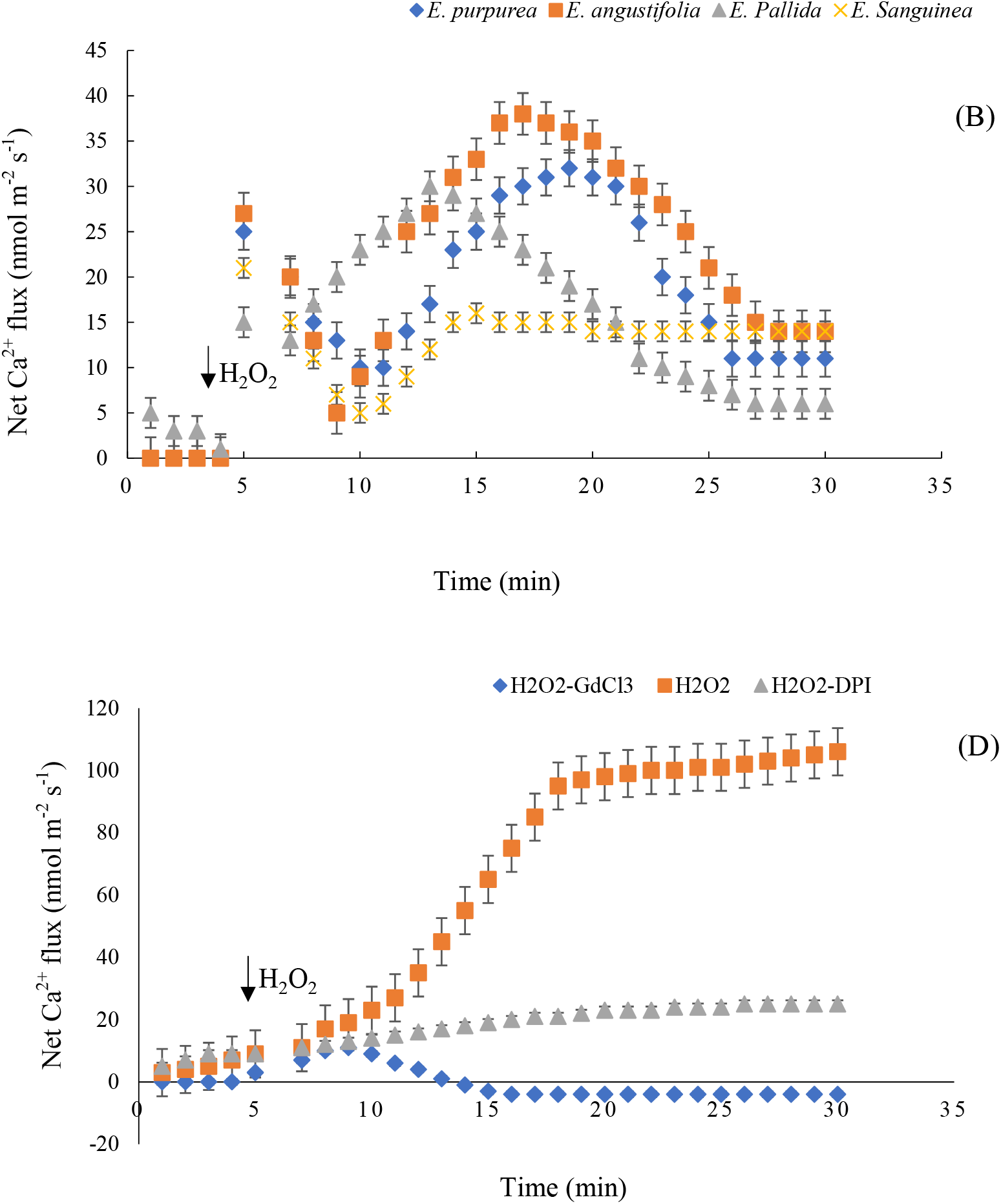
Transiently measured net Ca^2+^ flux responses of four *Echinacea* species measured from root elongation (EZ; A) and mature (MZ; B) zones exposed to 10 mM H_2_O_2_ stress. Total Ca^2+^ flux (C) measured from EZ and MZ was calculated over 20 min after the addition of 10 mM H_2_O_2_. Transient net Ca^2+^ flux (D) response to 10 mM H_2_O_2_ stress measured from EZ in the presence of inhibitors: 20 µM diphenyleneiodonium (DPI), NADPH oxidase blocker; 0.1 mM gadolinium chloride (GdCl_3_), known as non-selective cation channel (NSCC) blocker. Different letters indicate the significant difference at * P < 0.05 among different *Echinacea* species. The error bars indicate the standard error (SE) for all the replicated data for each treatment. Data are shown as mean ± SE (n = 5).

### Hydroxyl radical-induced ion flux responses in roots

Demidchik et al., (2010) reported that the chemical interaction of H_2_O_2_ with iron (Fe) and copper (Cu) caused form the hydroxyl radicals as one of the most aggressive forms of ROS in the root system. This interaction could affect ion homeostasis significantly (Kohler et al., 2003). The net Ca^2+^ uptake of different *Echinacea* species in the presence of 0.3 mM Cu/Ascorbate (Cu/Asc) in both root zones is shown in **Figure 5**. It was found that 0.3 mM Cu/Ascorbate (Cu/Asc) caused to increase in net Ca^2+^ uptake **(Figure 5)** in *Echinacea* species roots. Potassium flux was different in elongation and mature root zones between *Echinacea* species **(Figure 6)**, indicating the signaling role of hydroxyl radicals in plants.

**Figure 5.**
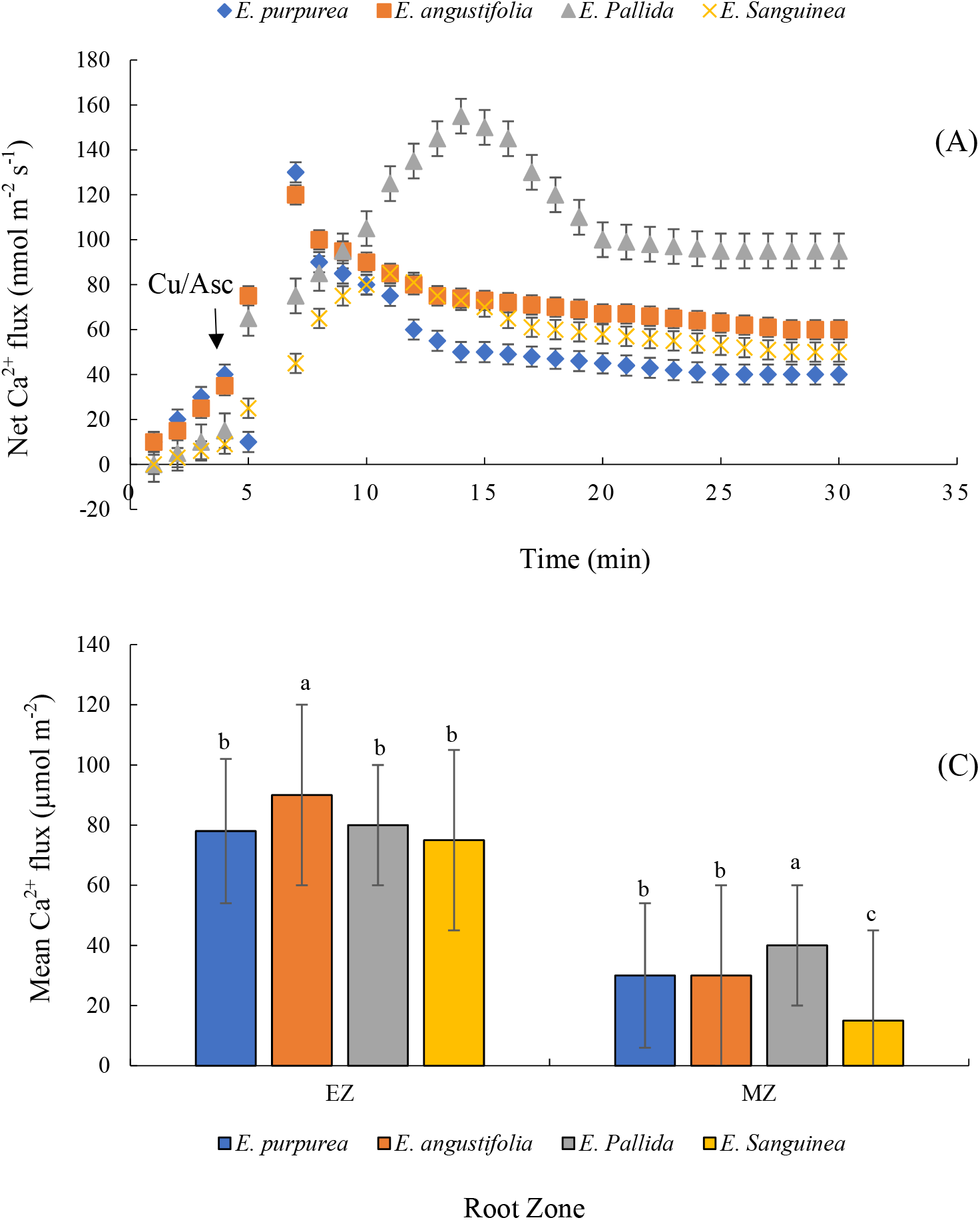
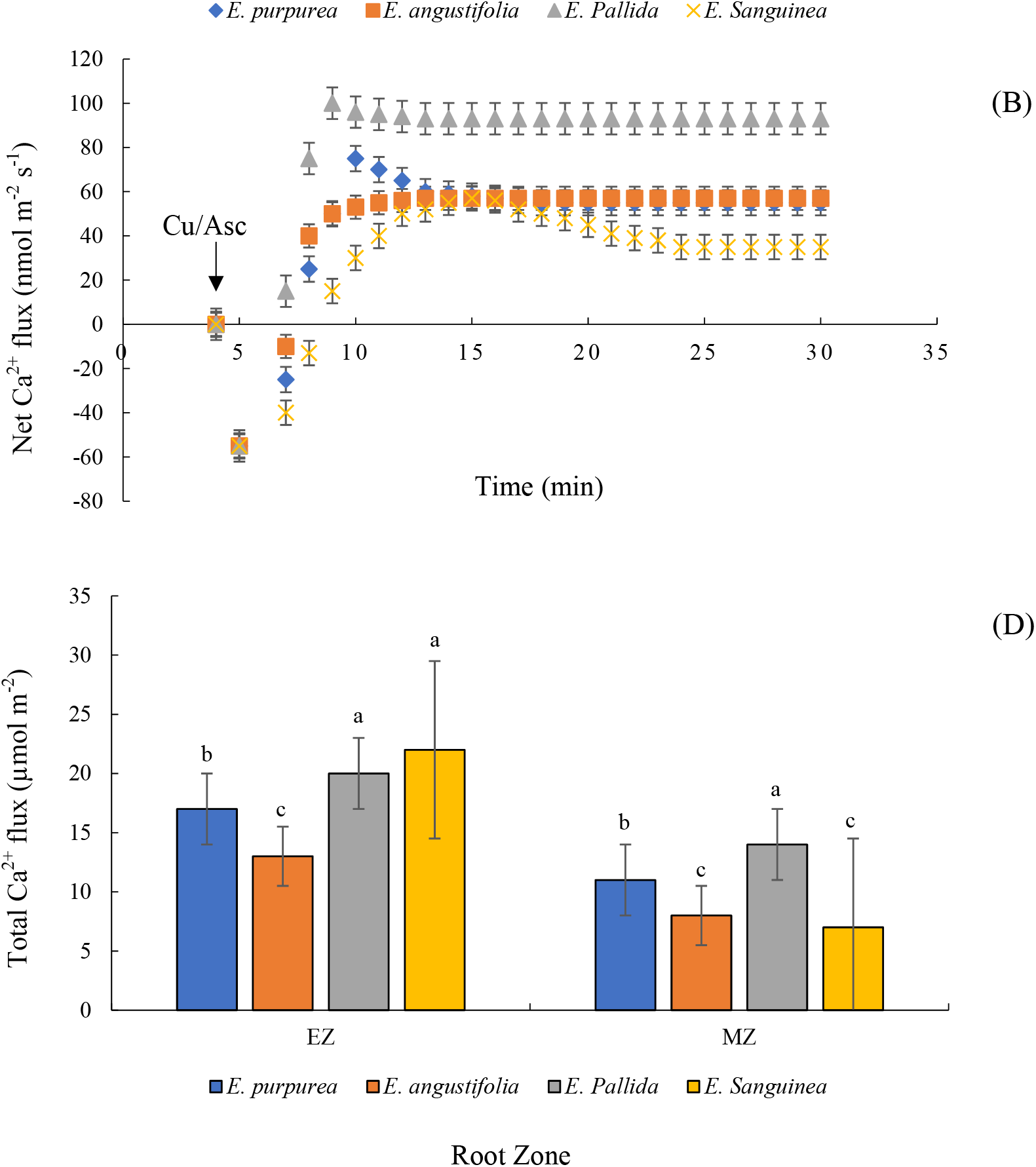
Transiently measured net Ca^2+^ flux response of four *Echinacea* species measured from the root elongation (A) and mature (B) zones exposed to 0.3/1 mM Cu/Asc stress. Mean and total Ca^2+^ (C-D) uptake were measured from the root elongation (EZ) and mature (MZ) zones. Mean and total Ca^2+^ fluxes were calculated over 20 min after the addition of 0.3/ 1 mM Cu/Asc. Different letters indicate the significant difference at * P < 0.05 among different *Echinacea* species within the same root zone. The error bars indicate the standard error (SE) for all the replicated data for each treatment. Data are shown as mean ± SE (n = 5).

**Figure 6.**
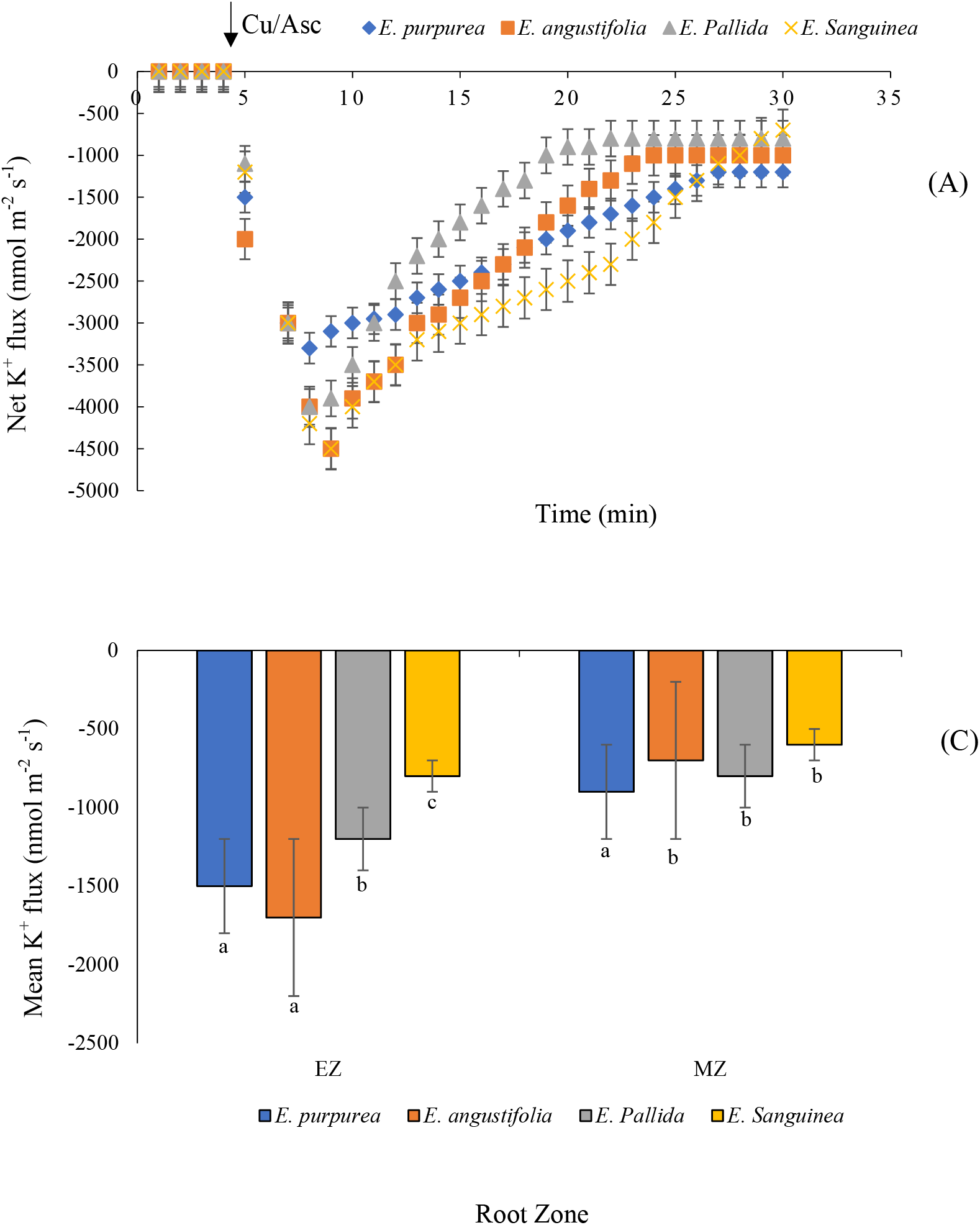
Transiently measured net K^+^ flux response of four *Echinacea* species measured from the root elongation (A) and mature (B) zones exposed to 0.3/1 mM Cu/Asc stress. Mean and total K^+^ lost (C-D) were measured from the root elongation (EZ) and mature (MZ) zones. Mean and total K^+^ fluxes were calculated over 20 min after the addition of 0.3/1 mM Cu/Asc. Different letters indicate the significant difference at * P < 0.05 among different *Echinacea* species within the same root zone. The error bars indicate the standard error (SE) for all the replicated data for each treatment. Data are shown as mean ± SE (n = 5).

### DHAR and MDHAR activities

The DHAR and MDHAR activities as affected by salinity stress in four *Echinacea* species are shown in **Figure 7AB**. Accordingly, the DHAR activity was decreased in *E. purpurea, E. angustifolia*, and *E. sanguinea* due to the salinity treatment **(Figure 7A)**. According to results (**Figure 7A**), 12.2% increase in DHAR activity was found in *E. pallida* on day 6 of salinity treatment, although a decrease was observed in the species **(Figure 7A)**. Meanwhile, the MDHAR activity was decreased in *all Echinacea* species toward the end of the experiment **(Figure 7B)**. The MDHAR activity decreased and was constant in the *E. purpurea* and *E. sanguinea* and *E. angustifolia* and *E. pallida* species, respectively on 6th and 12th days (**Figure 7B)**.

**Figure 7.**
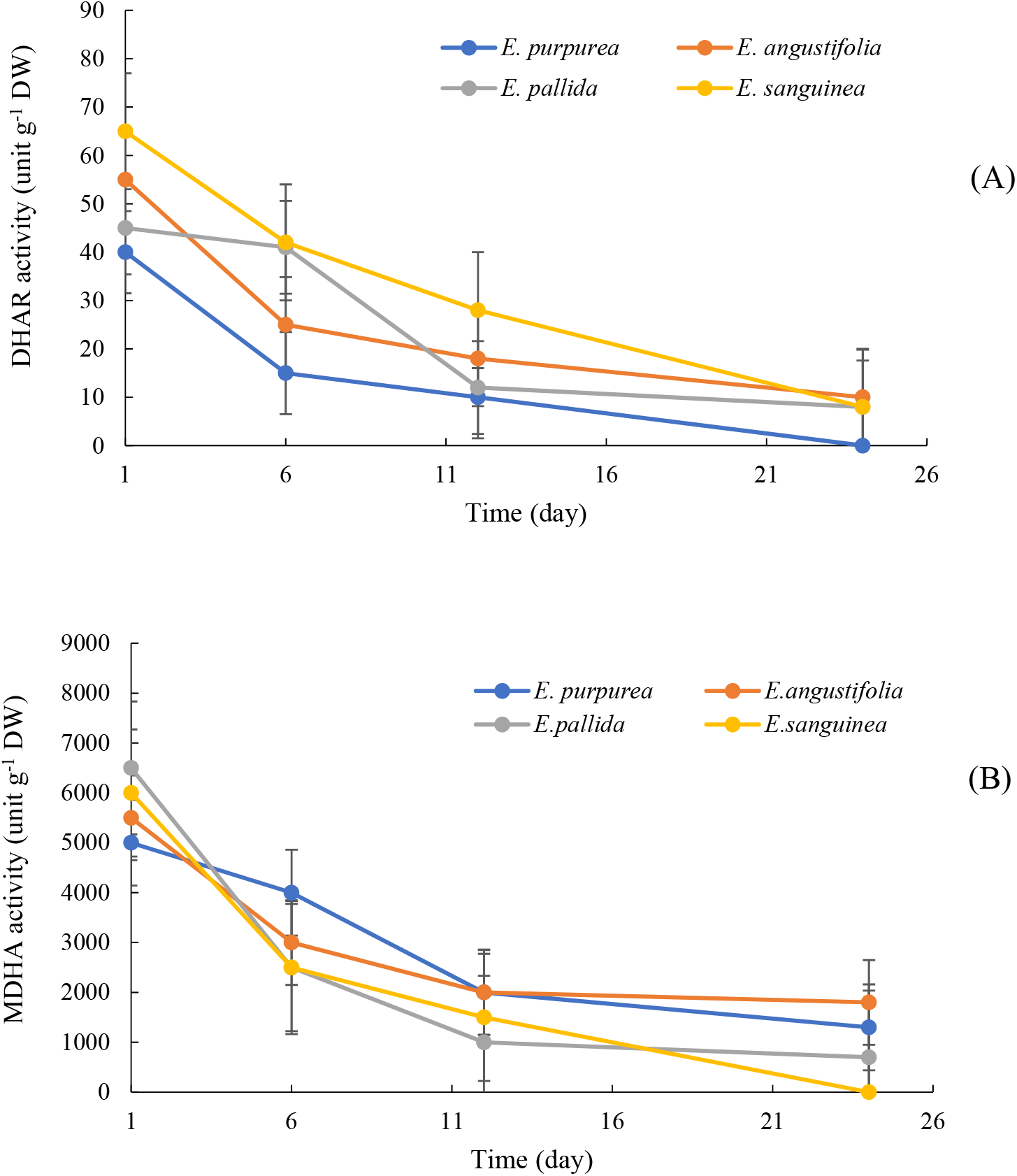
Dehydroascorbate reductase (DHAR) and monodehydroascorbate reductase (MDHAR) of four Echinacea species during the time. Vertical bars are the mean values ± standard errors (n = 5).

## DISCUSSION

### How K^+^ retention influence the *Echinacea* salinity tolerance?

Increasing root K^+^ efflux by increasing salinity stress, and plants’ sensitivity to salinity are reported in different plants (Sun et al., 2015; Liu et al., 2019). Anschütz et al., (2014) demonstrated the positive effect of cytosolic K^+^ levels in plants’ response to salinity stress. However, the effect of stress-induced K^+^ loss on the *Echinacea* species salinity tolerance it is still unclear. According to the results obtained in this study, the *E. angustifolia* species had a smaller K^+^ efflux than *E. purpurea* in the mature zone **(Figure 1BC)**, however, no clear pattern was found in the elongation region **(Figure 1AC)**. Potassium efflux was different between salt-sensitive and tolerant species, as it was 3-fold higher in *E. sanguinea* and *E. pallida* species than *in E. purpurea* and *E. angustifolia* **(Figure 1AB)**. So, it is possible that another pathway is involved to mediate the salinity stress in *E. sanguinea* and *E. pallida* species.

The main impacts of salinity stress are reducing plant growth, disturbing the ionic balance, and accumulation of oxygen species in plant tissues (Solis et al., 2021). The oxidative damage of membranes, chloroplasts, and mitochondria consequently due to ROS accumulation, cause to plant death immediately (De Pinto et al., 2012; Hossain et al., 2015). However, signaling role of ROS in stress conditions and regulator of ions-permeable channels were reported by Baxter et al., (2013) and Demidchik (2018).

The results of the present study demonstrated the positive influence of 10 mM H_2_O_2_ treatment on the K^+^ efflux in both root zones, especially in the elongation root zone **(Figure 3AC)**, demonstrate the function of K^+^ efflux as signaling factor in plant responses to salinity stress. Generally, various findings of the present study relevant that the adaptation strategies to salinity stress is depending on *Echinacea* species, and stress-induced K^+^ efflux plays a critical role in the determination of salinity tolerance in the cell-specific context.

### Calcium uptake as affected by H_2_O_2_ and ROS

Hydrogen proxied acts as a signaling molecule between plant organs and distant tissues (Baxter et al., 2013). Meanwhile, it is reported that H_2_O_2_ is related to Ca^2+^ signals (Steinhorst and Kudla, 2013; Gilroy et al., 2014). Bose et al., (2014) demonstrated that the H_2_O_2_ levels significantly increase as affected by salinity stress. So, in order to assess the H_2_O_2_ role on Ca^2+^ signals, 10 mM H_2_O_2_ was applied in the present study. Obtained results demonstrated that 10 mM H_2_O_2_ stimulated Ca^2+^ influx in all *Echinacea* species in the mature and elongation root zones, however, the *E. angustifolia* and *E. purpurea* species had more ROS-induced Ca^2+^ influx than *E. sanguinea* and *E. pallida* **(Figure 4AB)**.

Calcium influx as affected by ROS has been observed in sensitive species, as well, but it was two-fold higher in salt-tolerant species. The same results were found by Zhu et al., (2017). It again suggests the different strategies of salt-sensitive and tolerant *Echinacea* species. Reactive oxygen species can produce by NADPH oxidase, which is the main factor in net root Ca^2+^ uptake (Demidchik and Maathuis, 2007). Based on the results **(Figure 4D)**, only a 52% reduction was found in Ca^2+^ fluxes by NADPH oxidase activity and DPI in salt-tolerant *E. angustifolia*. So, it demonstrated the strong influence of NADPH oxidase on root Ca^2+^ signaling under salinity stress conditions. NADPH oxidase can be active by cytosolic Ca^2+^ that amplifies both ROS and Ca^2+^ signals (Demidchik et al., 2018).

### Voltage-gated channels in *Echinacea* species

**Figure 2AB** showed the significant increase in H^+^ pumping as affected by NaCl stress in both root zones. Orthovanadate had an important role in H^+^-ATPase in this process. The important point of this is the depolarization-activated GORK channel, which reduces K^+^ loss subsequently and restores the membrane potential (Wu et al., 2018). The ion fluxes plotting of H^+^ and K^+^ fluxes for four *Echinacea* species in the presence of salt stress is shown in **Figure 8**. Different *Echinacea* species groups were different based on the H^+^-ATPase activity and K^+^ efflux. lower K^+^ efflux was found in the *E. purpurea* and *E. angustifolia* group, but it was higher in H^+^ efflux, demonstrating that these species may remove Na^+^ ions by activated H^+^-ATPase pumping. Whereas, that was not found for *E. pallida* and *E. sanguinea* species. Once again, obtained results show the different strategies by four *Echinacea* species in salinity stress conditions.

**Figure 8.**
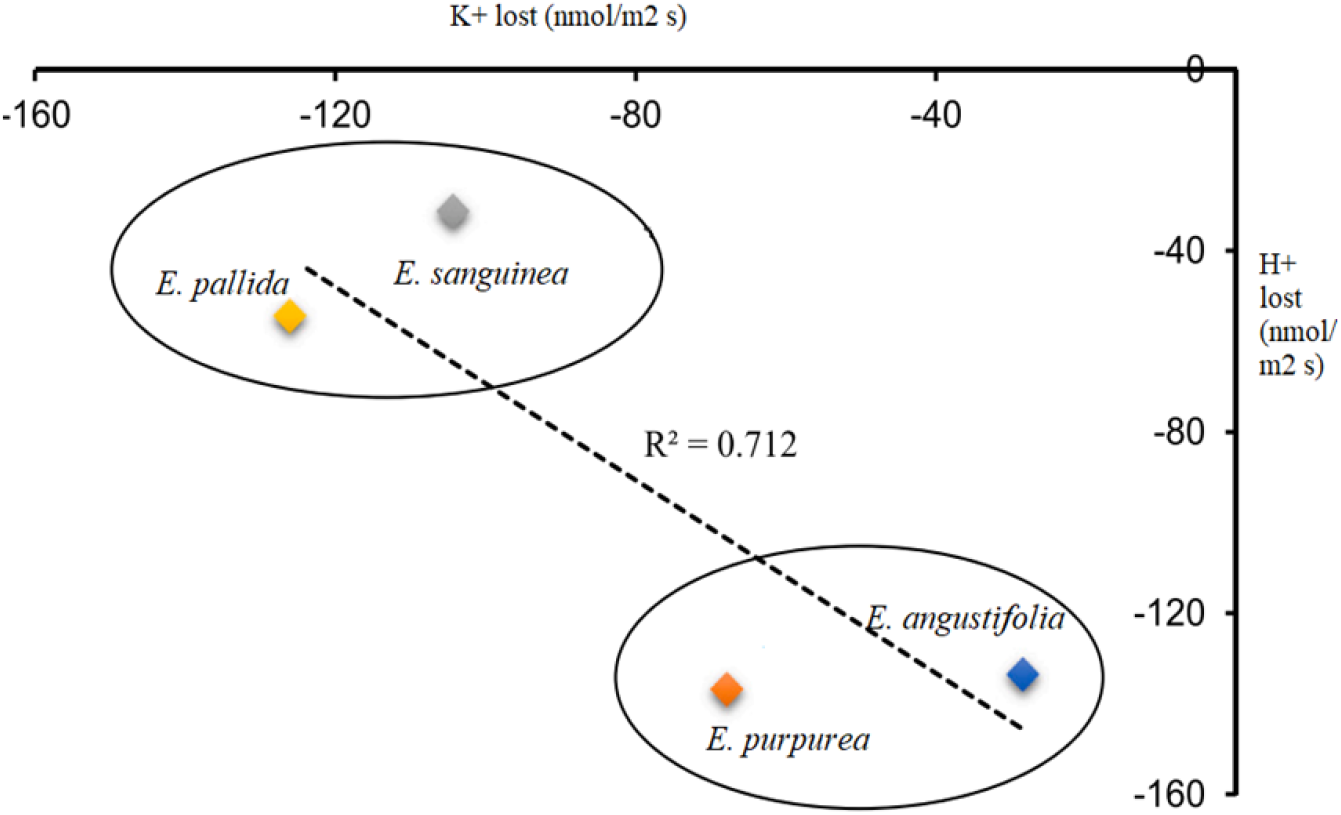
Relationship of H^+^ and K^+^ efflux in salinity tolerance in *Echinacea* species. Steady-state H^+^ and K^+^ fluxes were measured from the root mature zone of four *Echinacea* species calculated over 20 min after the addition of 100 mM NaCl.

In summary, the NaCl and ROS-induced K^+^ efflux was remarkably higher in the *E. pallida* and *E. sanguinea* species compared with *E. purpurea* and *E. angustifolia* species in the elongation root zone. Unlike the lower K^+^ efflux in the *E. purpurea* and *E. angustifolia* species, higher H^+^ efflux was found in these species caused by NaCl and ROS treatments. According to pharmacological results, TEA and GdCl_3_ significantly inhibited the K^+^ efflux as a result of NaCl and ROS, indicating the effective role of GORK and NSCC channels in K^+^ efflux. The significant effect of H^+^-ATPase in H^+^ efflux was demonstrated by Sodium orthovanadate. Generally, *E. purpurea* and *E. angustifolia* species were sensitive to NaCl and ROS in the elongation zone. The metabolic switch role of K^+^ efflux to adaptation with salinity stress is possible in the elongation root zone, which may provide an advantage to *E. pallida* and *E. sanguinea* in salinity stress conditions.

## MATERIALS AND METHODS

### Plant cultivation

*Echinacea purpurea, Echinacea angustifolia, Echinacea pallida*, and *Echinacea sanguinea* seeds were cultivated in 2.5-liter pots after sanitized with 10% bleach (10 min) and water-rinsed. Pots were filled with loamy soil texture. The experiment was carried out in greenhouse with average temperature 21/19^?^C (day/night), and 60% humidity. All soil physiochemical properties were determined based on the standard instructions described by Rowell (1994). The soil samples were neutral in pH (average 7.6), non-saline (average EC 1.65 dS m^-1^), and calcareous (average 15.6% CCE) in nature. Total N, available -P, available – K, and soil OM were 0.28%, 18.25 mg kg^-1^, 215 mg kg^-1^, and 0.87% in average, respectively. Plants were irrigated every 6 h with saline water (100 mM NaCl) for five months to avoid osmotic shock and harvested after eight months. The roots with a defined length (approximately 10-15 cm) were kept for further electrophysiological analyses.

### Non-invasive ion flux

The non-invasive MIFE approach was performed to measure the ions (Ca^2+^, K^+^, and H^+^) fluxes (Shabala et al., 1997). Briefly, the different *Echinacea* species’ roots were immobilized in the Petri dish containing BSM solution with 0.5 mM KCl and 0.1 mM CaCl_2_. The Petri dishes were kept at 25 ± 1^?^C for 1h. Two root zones, elongation and mature area, were assessed for ion fluxes. Three treatments including 0.3/1 mM CuCl_2_/Ascorbate (Cu/Asc), 10 mM H_2_O_2_, and 100 mM NaCl were performed for steady-state fluxes (Shabala et al., 2022). The CHART program and Nernst slope of the electrode was applied for different ion fluxes, and Net fluxes were calculated using MIFEFLUX software (Shabala et al., 1997).

### Pharmacological experiment

Different kinds of blockers of H^+^-ATPase, K^+^-selective channel blockers, the blocker of NADPH oxidase, and a blocker of non-selective cation channels (NSCC), including 1 mM sodium orthovanadate (OV, Na_3_VO_4_), 20 mM tetraethylammonium chloride (TEA), 0.1 mM gadolinium chloride (GdCl_3_), and 20 µM diphenyleneiodonium (DPI), were applied in root *Echinacea* species for 1 h, respectively (Shabala et al., 2022).

### DHAR and MDHAR activities

50 mM potassium phosphate buffer (pH 7) was used to measure the dehydroascorbate reductase (DHAR) activity, mixed with 0.2 mM DHA, 0.1 mM EDTA, and 2.5 mM GSH in a final volume of 1 mL at 265 nm. (Nakano and Asada, 1981). The monodehydroascorbate reductase (MDHAR) activity was determined at 340 nm and absorbance coefficient of 6.2 mM (Hossain and Asada, 1984).

### Statistical analysis

Student’s t-test (P<0.05) was performed using Statistical analysis software (SAS, 9.4). Data shown as means ± standard error (SE).

## FUNDING INFORMATION

This research did not receive any specific grant from funding agencies in the public, commercial, or not-for-profit sectors.

